# Creating an atlas of the bone microenvironment during oral inflammatory-related bone disease using single-cell profiling

**DOI:** 10.1101/2022.09.13.507807

**Authors:** Yi Fan, Ping Lyu, Ruiye Bi, Chen Cui, Ruoshi Xu, Clifford J. Rosen, Quan Yuan, Chenchen Zhou

## Abstract

Oral inflammatory diseases such as apical periodontitis are common bacterial infectious diseases that may affect the periapical alveolar bone tissues. A protective healing process occurs simultaneously with the inflammatory tissue destruction, in which mesenchymal stem cells (MSCs) play a primary role. However, a systematic and precise description of the cellular and molecular composition of the microenvironment of bone affected by inflammation is lacking. In this study, we created a single cell atlas of cell populations that compose alveolar bone in healthy and inflammatory disease states. We investigated changes in expression frequency and patterns related to apical periodontitis, as well as the interactions between MSCs and immunocytes. Our results highlight an enhanced self-supporting network and osteogenic potential within MSCs during apical periodontitis-associated inflammation. MSCs not only differentiated towards osteoblast lineage cells, but also expressed higher levels of osteogenic related markers, including SPARC and Col1α1. This was confirmed by lineage tracing in transgenic mouse models and human samples from oral inflammatory-related alveolar bone lesions. In summary, the current study provides an in-depth description of the microenvironment of MSCs and immunocytes in both healthy and disease states. We also identified key apical periodontitis-associated MSC subclusters and their biomarkers, which could further our understanding of the healing process and the underlying mechanisms of oral inflammatory-related bone disease. Taken together, these results enhance our understanding of heterogeneity and cellular interactions of alveolar bone cells under pathogenic and inflammatory conditions. We provide these data as a tool for investigators not only to better appreciate the repertoire of progenitors that are stress responsive but importantly to help design new therapeutic targets to restore bone lesions caused by apical periodontitis and other inflammatory-related bone diseases.

## Introduction

Oral diseases, particularly dental caries and periodontal diseases, affect 3.5 billion people worldwide (Disease, Injury, & Prevalence, 2018). Untreated dental caries can directly lead to pulp necrosis and periapical lesions, resulting in apical periodontitis. Individuals with at least one tooth affected by apical periodontitis comprise up to 52% of cases, indicating that apical periodontitis is a highly prevalent disease (Tiburcio-Machado et al., 2021). Inflammation in the oral cavity can lead to destruction of surrounding periapical tissues and resorption of hard tissues, a consequence of the unbalanced interaction between infection and the immune response (Gazivoda et al., 2009; Márton & Kiss, 2014). Restoring and regenerating the destroyed periapical alveolar bone structures has always been a challenging task in clinical practice. Active inflammation, the tissue injury and the protective healing process all occur simultaneously in the setting of chronic apical periodontitis (Márton & Kiss, 2014). Importantly, there is a complex assemblage of immune cell types involved in the pathogenesis, highlighting the importance of polymorphonuclear leukocytes (PMN), lymphocytes and monocyte/macrophages in periapical defense (Braz-Silva et al., 2019; Nair, 2004). Notably, an increasing number of studies report the involvement of mesenchymal stem cells (MSCs) in the protective action that occurs during oral inflammatory diseases, whereby MSCs exert immunomodulatory effects and have regenerative potential (Li et al., 2014; Marton & Kiss, 2000; Nair, 2004). MSC markers such as CD44, CD73, CD90, CD106 and STRO-1 have been observed in human periapical inflammatory tissues (Estrela et al., 2019; J. Liao, Al Shahrani, Al-Habib, Tanaka, & Huang, 2011). Cells isolated from the inflamed periapical region were able to produce colony-forming unit-fibroblasts (CFU-Fs) with high osteogenic capacity. It is also reported that interference with MSC mobilization toward the periapex region in an apical periodontitis mouse model led to enlargement of lesions, accompanied by decreased wound healing markers and increased inflammatory cytokines (Araujo-Pires et al., 2014). These findings indicate the involvement of MSCs in the repair and regeneration of oral inflammatory-related bone lesions. They also suggest that MSCs present promising targets for treating bone lesions, with great potential for modulating inflammation and promoting tissue regeneration. However, most studies have surveyed whole tissues to understand the transcriptomic and cellular profile of these diseases. Specific cell populations and their regulatory molecules, as well as the interaction among different cell populations, remain far from clear. The advances in single-cell technologies offer an unbiased approach for identifying heterogeneous cell subsets, tracking the trajectories of distinct cell clusters and uncovering regulatory relationships between genes (Hwang, Lee, & Bang, 2018; Tang et al., 2009). In this study, we collected mandibular alveolar bone samples from control and apical periodontitis in mice and subjected them to scRNA sequencing (scRNA-seq). The atlas of the mandibular alveolar bone explored the distinct cell subsets and their expression profiles relevant to apical periodontitis. We also investigated the relationship between MSCs and immune cell subsets. The results reveal the role of a subset of MSCs in inflammation, which showed increased frequency and which formed a self-supporting network. Moreover, MSCs exhibited upregulated osteogenic potential, which was confirmed in transgenic mouse models and human patients with chronic apical periodontitis. These results advance our understanding of heterogeneity and interactions of alveolar bone cells in the pathogenesis of inflammatory-related bone diseases. Defining key cellular subsets such as MSCs and their biomarkers in inflamed tissue will be important for identifying new therapeutic targets for oral inflammatory-related bone diseases.

## Results

### Single cell transcriptional profiling identified 15 discrete populations in homeostasis and chronic apical periodontitis samples

Individual cells were isolated from alveolar bone of healthy mice and mice with apical periodontitis (AP). We modeled AP using a well-established AP mouse model in which the mandibular first molar pulp was exposed and subsequently developed chronic AP over a 3-week period (Taira et al., 2019). Bar-coded cDNA libraries from individual cells were obtained using the 10X Genomics Chromium Controller platform (Zheng et al., 2017) (Figure 1A). The combined libraries from healthy and AP alveolar bone contained 15148 individual cells. The median value of feature_RNA was between 1000 and 2000 (Figure 1-figure supplement 1B). After quality control filtering and removal of the batch effect between batches, the t-stochastic neighbor embedding (t-SNE) method was applied to reduce the dimensionality. Seurat’s unbiased cluster detection algorithm identified 15 distinct cell populations (Figure 1B, C). Cluster-specific transcripts were utilized to annotate cell types with classic markers as described in a previous study (Lin et al., 2021). These included B cell (CD79a), hematopoietic stem cell (HSC) (Cd34), Mesenchymal stromal cell (MSC) (Col1α1), natural killer (NK) cell (Klrd1), T cell (Cd3g), dendritic cell (Siglech), epithelial cell (Epcam), erythrocyte (Hbb-bt), macrophage (Adgre1), mast cell (Fcer1a), megakaryocyte (Gp1bb), monocyte (Ly62c), myeloid progenitor (Mpo), neutrophil (S100a8), pre-B cell (Vpreb1) (Figure 1E, F). The top 20 enriched genes in each defined cluster were identified and compared (Figure 1G).

**Figure 1.**
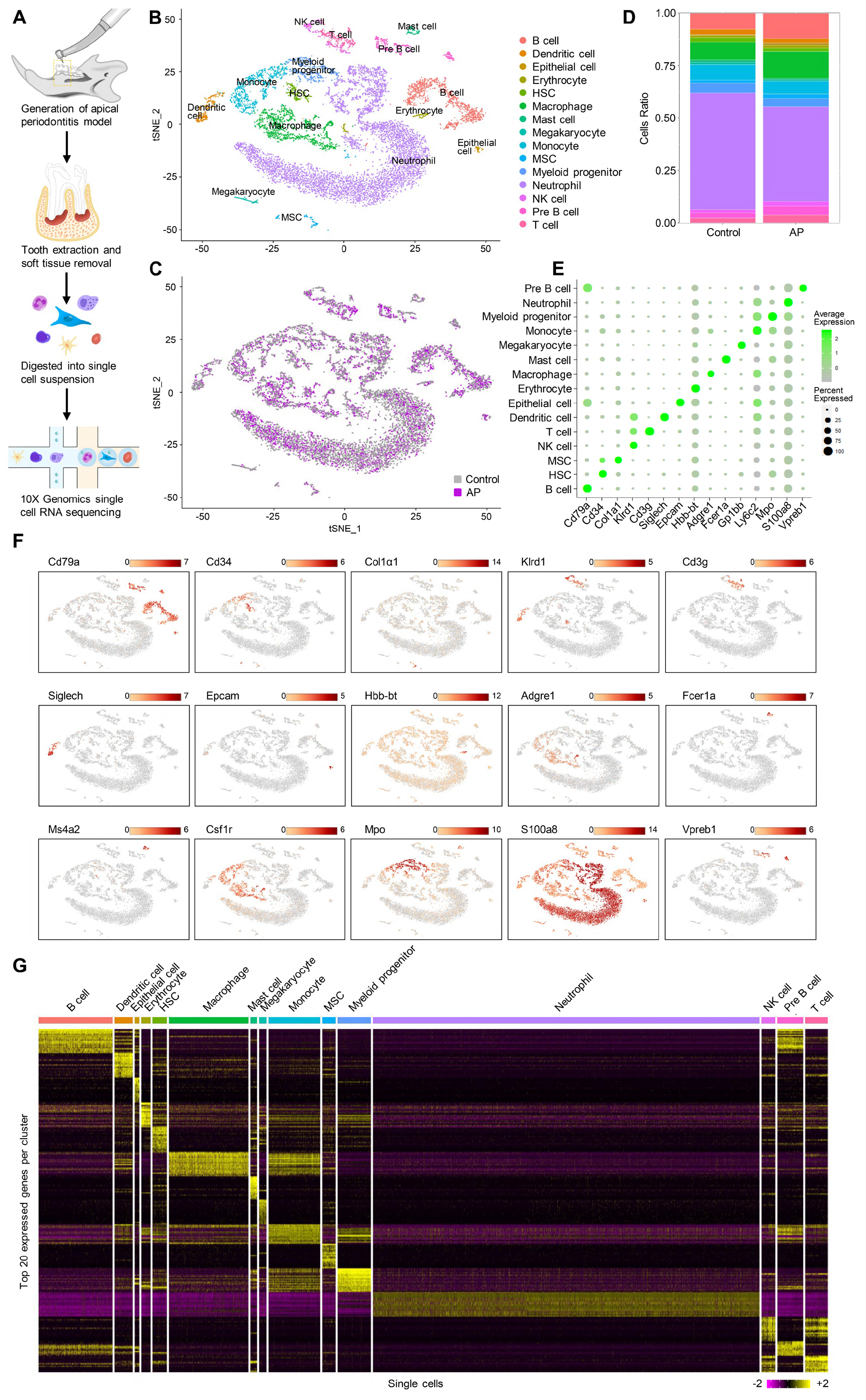
Identification of the single-cell atlas of alveolar bone using scRNA-seq and unbiased clustering. (A) Schematic diagram of the experimental design. (B-C) t-Stochastic neighbor embedding (t-SNE) representation of aligned gene expression data in single cells extracted from mandibles of control mice (n=8340) and apical periodontitis mice (n=6808) showing 15 distinct clusters and cellular origin. (D) Relative abundance of 15 cell populations composing alveolar bone under healthy and AP conditions. (E) Expression of gene markers in distinct cell types. (F) Gene expression patterns projected onto t-SNE plots of marker genes. scale: log-transformed gene expression. (G) Heatmap showing the 20 most upregulated genes (ordered by decreasing Padj value) in each cluster defined in B. scale: log2 fold change.

### AP led to significant changes in frequency and transcriptional expression of cell populations

All the identified cell clusters were present in both AP and control samples, but there were significant differences in the cellular compositions of particular clusters. T cell, B cell, NK cell, macrophage, epithelial cell and MSC had significantly increased frequency in AP samples. Neutrophil, myeloid progenitor, monocyte, megakaryocyte, mast cell, HSC and dendritic cell were markedly decreased (Figure 2A and 1D).

**Figure 2.**
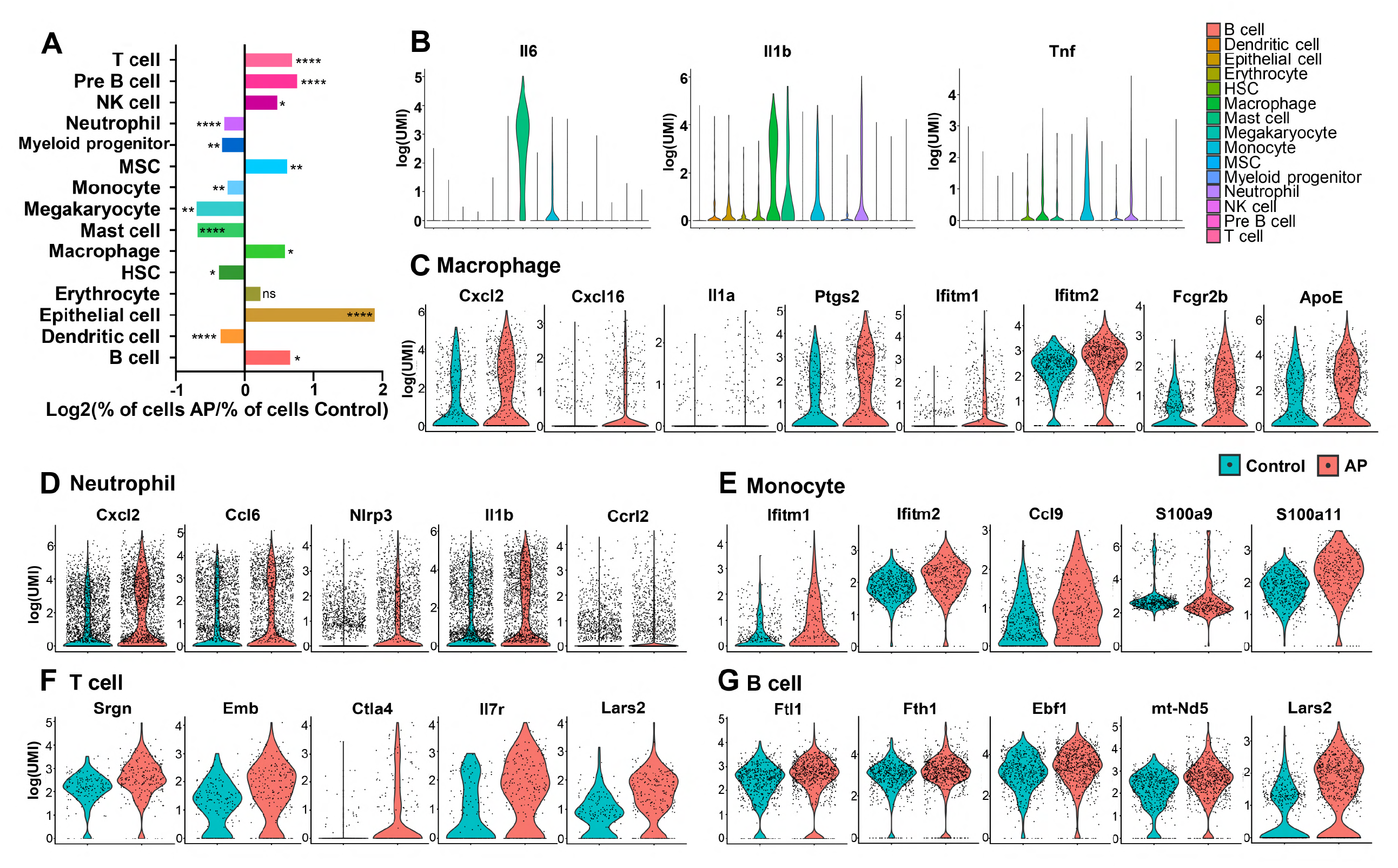
Changes in frequency and transcriptional expression pattern in each cell population from control and AP groups. (A) Bar plot of cells per cluster (AP versus Control). Normalization to overall number of inputs per condition. Fisher’s exact test with Bonferroni correction was used. (B) Violin plots of cluster-specific expression of representative genes. (C-G) Violin plots showing genes that significantly changed in each cluster from control and AP.

AP is a complex inflammatory process involving innate and adaptive immune responses (Cotti, Schirru, Acquas, & Usai, 2014). A variety of inflammatory cells such as neutrophils, mast cells, monocytes, macrophages and lymphocytes are involved in periapical lesions, highlighting the direct involvement of the immune response in the pathogenesis of AP (Nair, 2004). Neutrophils are important components in the acute phase of AP as a first line of defense. But they are also important in the progression of AP by interacting with microorganisms, leading to tissue damage and chemotaxis (Braz-Silva et al., 2019). Single-cell differential expression analysis revealed that the most significantly enriched genes in neutrophils were various proinflammatory chemokines and cytokines. These included C-X-C motif chemokine ligand 2 (Cxcl2), C-C motif chemokine ligand 6 (Ccl6), NLR family pyrin domain containing 3 (Nlrp3) and Interleukin-1β (IL-1β). Notably, we found that C-C motif chemokine receptor like 2 (Ccrl2) is upregulated in neutrophils during AP (Figure 2D). It is responsible for the innate defense against pathogens and is also involved in the regulation of neutrophil migration (Del Prete et al., 2017; Kolaczkowska & Kubes, 2013; Mantovani, Cassatella, Costantini, & Jaillon, 2011).

Mast cells, monocytes and macrophages have critical roles in the inflammatory infiltrate during chronic AP (Braz-Silva et al., 2019). The production of Interleukin-6 (IL-6) was present in these cell populations with the highest expression level in mast cells. The pro-inflammatory cytokine IL-1β is a key mediator of host response to microbial infection and is associated with the persistence of AP (Morsani, Aminoshariae, Han, Montagnese, & Mickel, 2011; Ng, Mann, Rahbaran, Lewsey, & Gulabivala, 2008). We found IL-1β transcripts in a series of cell types, such as monocyte, macrophage, mast cell and neutrophil. Of these, macrophages had the highest IL-1β expression. Another major cytokine, tumor necrosis factor (TNF) (Cotti et al., 2014), was detected in immunoresponsive cell clusters, such as monocyte, macrophage, mast cell, myeloid progenitor, neutrophil and HSC, with the highest expression observed in the monocyte population (Figure 2B).

Furthermore, gene signatures from monocytes showed that the interferon-induced transmembrane (IFITMs) protein 1 and 2 (Ifitm1 and Ifitm2) (Figure 2E) were upregulated the most during AP. These factors have been associated with signal transduction of anti-inflammation activity in the immune system (Yanez, Ross, & Crompton, 2020). We also detected upregulated expression levels of Ccl9 in the monocyte population from AP. Ccl9 is an important cytokine and is involved in the survival of osteoclasts during the destruction of the periapical bone (Silva, Garlet, Fukada, Silva, & Cunha, 2007). Also, genes coding for pro-inflammatory calcium-binding S100 family proteins such as S100a9 and S100a11 had increased expression in the AP monocyte cluster (Figure 2E). Previous reports demonstrated that macrophages are capable of secreting pro- and anti-inflammatory substances which act on the development and repair of the AP lesions (Italiani & Boraschi, 2014; Shapouri-Moghaddam et al., 2018). Indeed, several genes encoding pro-inflammatory mediators, including Cxcl2, Cxcl16, IL-1α, Ptgs2 were upregulated in macrophages from AP samples (Figure 2C). Expression of anti-inflammatory-associated genes such as Ifitm1 and Ifitm2 was significantly increased in the AP macrophage cluster compared to control cells. Furthermore, Fcγ receptor IIB (Fcγr2b) was markedly upregulated in macrophages from AP samples. Fcgr2b is expressed in most tissue-resident macrophages (Gautier et al., 2012) and functions to inhibit Fcγ-dependent phagocytosis. It also inhibits release of cytokines such as IL-6, TNF-α, IL-1α, as well as neutrophil chemotactants (Clatworthy & Smith, 2004; Espeli, Smith, & Clatworthy, 2016). In addition, expression of Apolipoprotein E (ApoE), which can suppress the pro-inflammatory response (Jofre-Monseny et al., 2007), was significantly increased in the macrophage population (Figure 2C). These data indicated the activation of anti-inflammatory factors by macrophages during local inflammation by AP.

The major classes of lymphocytes are T lymphocytes, B lymphocytes and the NK cells. T and B lymphocytes comprise the majority of the inflammatory infiltrate in AP (Graunaite, Lodiene, & Maciulskiene, 2012). A significant increase in the expression of inflammatory-associated genes, such as Srgn, Emb, Ctla4 and Il7r could be observed in the AP T cell population (Figure 2F). In the AP B lymphocytes cluster, inflammation-responsive genes (Fth1, Ftl1, Ebf1, mt-Nd5) were upregulated (Figure 2G). Interestingly, Lars2, the gene encoding a mitochondrial leucyl tRNA synthase (Carminho-Rodrigues et al., 2020; t Hart et al., 2005), was significantly upregulated in T and B lymphocytes, indicating changes in mitochondrial metabolism in both clusters.

### AP leads to reduced transcriptionally inferred cellular interactions with an increased self-supporting network in MSCs

We next sought to characterize the cell-cell communication related to the perturbation of signaling pathways detected in the AP samples by employing CellphoneDB (Efremova, Vento-Tormo, Teichmann, & Vento-Tormo, 2020; Nagai, Leimkuhler, Schaub, Schneider, & Costa, 2021). We identified a close interaction among MSC, macrophage and dendritic cells under homeostasis conditions (Figure 3A). A similarly close communication was found among MSC, macrophage and dendritic cells under inflammatory conditions (Figure 3B). Next, we compared the differential cell-cell interaction (CCI) network between AP and control samples using CrossTalkeR (Nagai, Leimkuhler, et al., 2021). The results suggested that AP is associated with an overall decrease in cellular interactions. However, MSC intercellular communication with mast cells and dendritic cells is upregulated and accompanied by the highest number of interactions within MSCs themselves (Figure 3C). These results indicate that, although cell populations lose their normal physiological interactions, MSCs were able to establish a self-interacting network and coordinate with certain types of cells during chronic AP. Next, we ranked the individual ligands by the number of their interactions. Inflammatory-related proteins (*Lgals9, Tnf, Ccl4*), extracellular matrix protein (*Fn1*), protein involved in biomineralization (*Spp1*) were among the highest interactions. Also, Tgfb1, Vegfb and Vegfa were in the top ten most abundant ligands (Figure 3D). Bar plots were also generated to display the top ten upregulated gene/cell pairs, showing that inflammation associated genes (Tnf/neutrophil, Ccl3/mast cell, Ccl3/monocyte, Il1β/macrophage) and matrix related genes (Sele/MSC, Fn1/MSC, Fn1/monocyte) were the most influential ligands during AP when compared to control state (Figure 3E). We used a Sankey plot to further focus on MSC-related interactions (Figure 3F). The results indicated that Sele was primarily directed by MSC towards the MSC cluster via multiple receptors including Glg1, Selplg and CD44. Moreover, Fn1 was secreted by MSC, monocyte, macrophage and mast cells towards MSC cells (Figure 3G). Of note, Sele and Fn1 are important cell adhesion molecules that mediate cell homing and migration (Frenette, Subbarao, Mazo, von Andrian, & Wagner, 1998; To & Midwood, 2011). This was in accordance with the increased frequency of the MSC cluster (Figure 2A) as well as the upregulated cellular interaction among MSCs and other cell populations.

**Figure 3.**
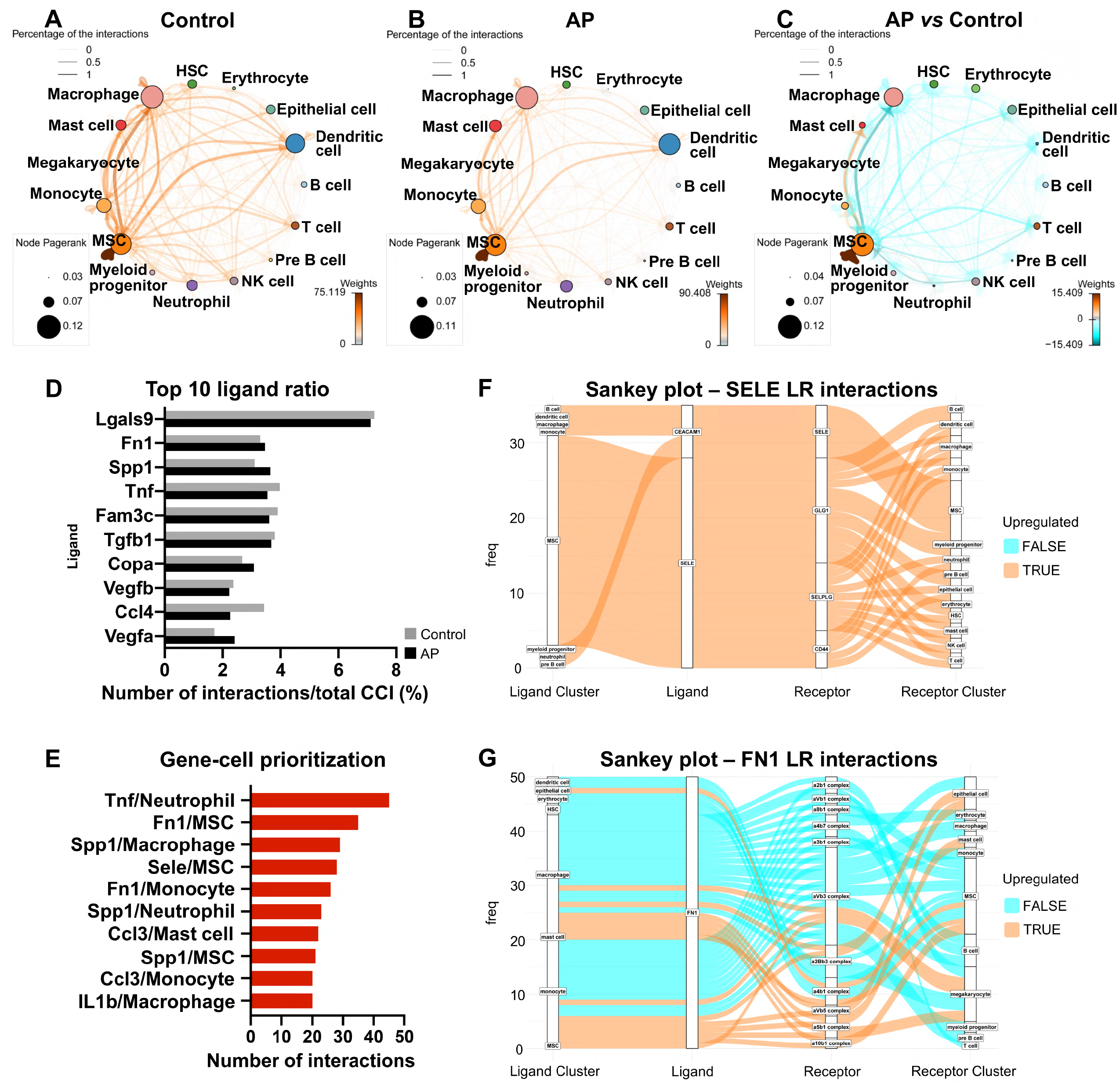
AP suppressed transcriptionally inferred cellular interactions and increased a self-supporting network within the MSC cluster. (A-C) Network plot of ligand-receptor activity in control (A), AP (B) and AP versus control (C). (D) Bar plot of top ten most abundant ligands in all inferred ligand-receptor interactions. (E) Ranking of Ligand/Source regarding communication gains in AP state. (F, G) Sankey plot listing all predicted source, receptor and receiver interactions associated with Fn1 and Sele.

### scRNA-seq based identification of AP-associated MSC population

MSCs represented the non-immune cell population in alveolar bone, constituting 1.76% of total identified cells. This cell population could be decomposed into four subclusters (Figure 4A). The most dominant subcluster was characterized by high expression of Prrx1, platelet-derived growth factor receptor (PDGFR)-β, and hematopoiesis supporting factors such as C-X-C motif chemokine 12 (Cxcl12), angiopoietin (Angpt1). It was also characterized by osteogenic-related markers, such as Runt-related transcription factor 2 (Runx2), Sp7, and was thus classified as the MSC_osteolineage subcluster (OLCs). The other subclusters were identified as MSC_endothelial (Cdh5^+^), MSC_neurological (Plp1^+^) and MSC_inflammatory (S100a8/S100a9^+^) (Figure 4A, B). The cell composition of the MSC cluster indicated an expansion of MSC_inflammatory subcluster during AP, whereas the MSC_neurological subcluster was reduced (Figure 4C). Of note, we identified increased numbers of Prx1^+^ and Sp7^+^ MSC_OLCs in AP lesions (Figure 4-figure supplement 1A).

**Figure 4.**
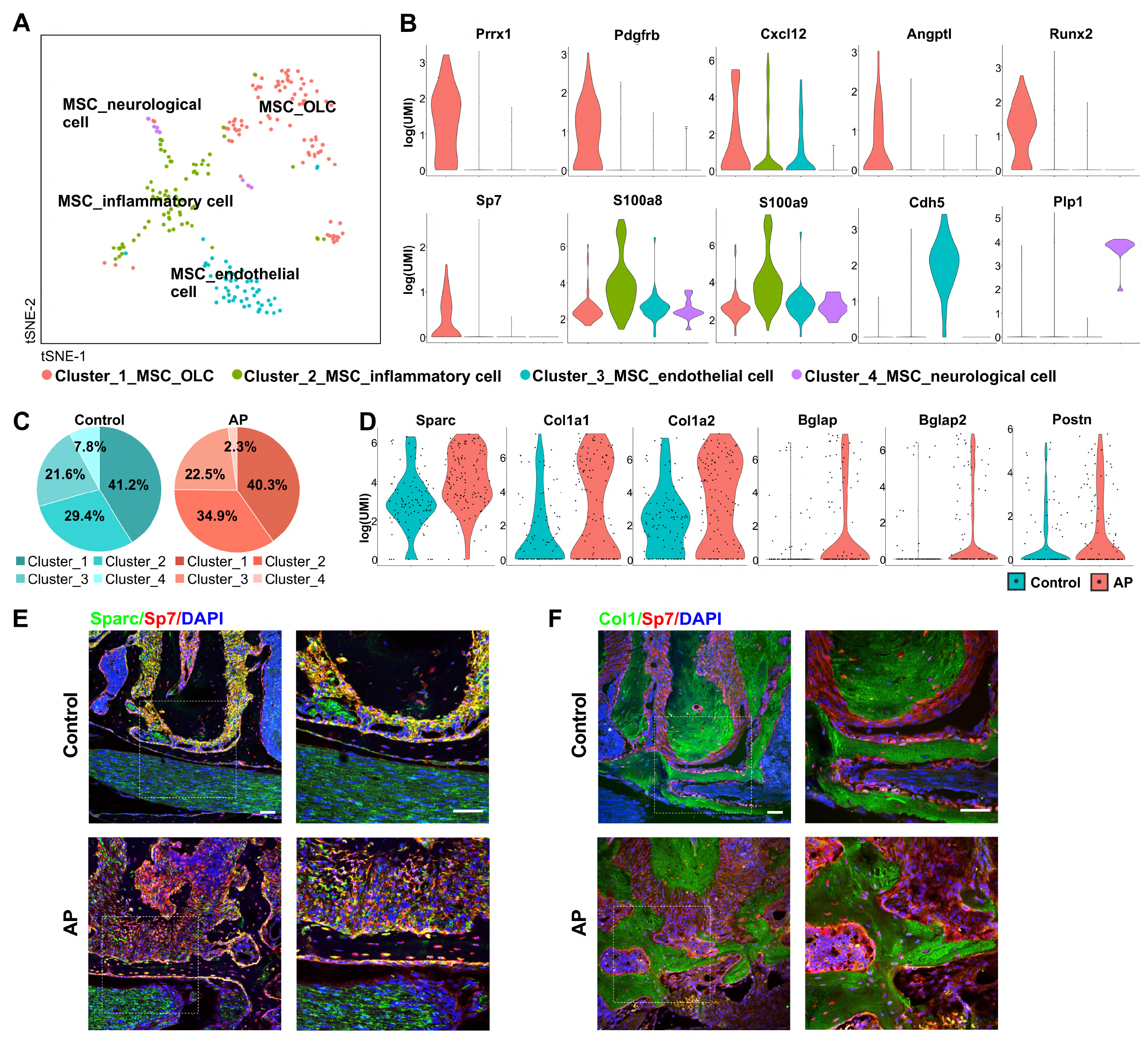
Identification and characterization of AP-associated MSC population and its subclusters. (A) t-SNE representation an unsupervised clustering of single cells within the MSC cluster. (B) Violin plots of MSC subcluster-specific expression of representative genes. (C) The percentages of four subclusters of MSC population were quantified in control and AP groups. (D) Violin plots depict the changes in the expression of top upregulated genes in the MSC cluster. (E, F) Immunofluorescence double staining of Sparc (E) and Col1 (F) in Sp7-expressing osteoprogenitors, n=6. Scale bar, 50μm.

We compared the MSC marker genes, identified above, among the four subpopulations. Their respective Gene Ontology (GO) enrichment analysis confirmed that there are four specific MSC populations. MSC_OLCs were enriched for ossification, connective tissue development, skeletal system development and osteoblast differentiation. MSC_endothelial cells displayed enrichment for angiogenesis, blood vessel morphogenesis and endothelium development. MSC_neurological cells were enriched for myelination, ensheathment of neurons and axon ensheathment. GO term analyses suggested highly specialized functional features of MSC_inflammatory cells, including regulation of protein export from nucleus, electron transport chain, mitochondrial ATP synthesis coupled electron transport and oxidative phosphorylation (Figure 4-figure supplement 2A).

### AP stimulates MSC differentiation towards osteoblast lineage cells

We next compared the transcriptome data from the MSC cluster between AP and control groups. Among the top upregulated genes, we observed a significant difference in the single cell expression levels of OLC-associated transcripts, such as secreted protein that is acidic and rich in cysteine (*Sparc), Col1α1, Col1α2, Bglap, Bglap2* and *Postn* (Figure 4D). *In vivo* experiments confirmed the upregulated expression of osteogenic-related markers, such as Sparc and Col1α1 in the AP group. There were increased numbers of Sparc^+^/Sp7^+^ or Col1^+^/Sp7^+^ cells embedded in the bone matrix, implying the protective function of MSC to differentiate into osteoprogenitors and osteoblasts under AP conditions (Figure. 4E, F).

We investigated the heterogeneity of MSCs by applying branch expression analysis modelling (BEAM) and corresponding pseudotime analysis implemented in Monocle 2 (Qiu, Hill, et al., 2017). The results revealed three distinct states. Of these, state 2 and state 3 represented two diverse differentiated cell populations (terminal branch) (Figure 5A). We then examined the characteristics of the MSCs with respect to their specific states. We plotted the estimated pseudotime of each cluster cell in the state space and the results indicate that starting with MSC_endothelial cells, MSCs were able to differentiate into MSC_OLCs and MSC_neurological cells. MSC_inflammatory cells may correspond to an intermediate state (Figure 5B). Interestingly, MSC_OLCs in state 3, as osteogenic primed MSCs, exhibited the highest tendency towards osteoblastic differentiation with upregulated expressions of Col1α1, Col1α2, Bglap, Spp1 and Postn (Figure 5C). Notably, this MSC population increased significantly in the AP group (Figure 5D). A comparison of the dynamics of the gene expression pattern across pseudotime revealed upregulation of osteogenic genes such as Sparc, Col1α1, Col1α2 and Bglap during inflammation when compared to the homeostatic state (Figure 5E). Thus, the MSC subcluster exhibited gene expression levels of osteogenic markers that increased during AP. These results were in accordance with our previous single cell analysis of gene expression in the AP group (Figure 4D-F).

**Figure 5.**
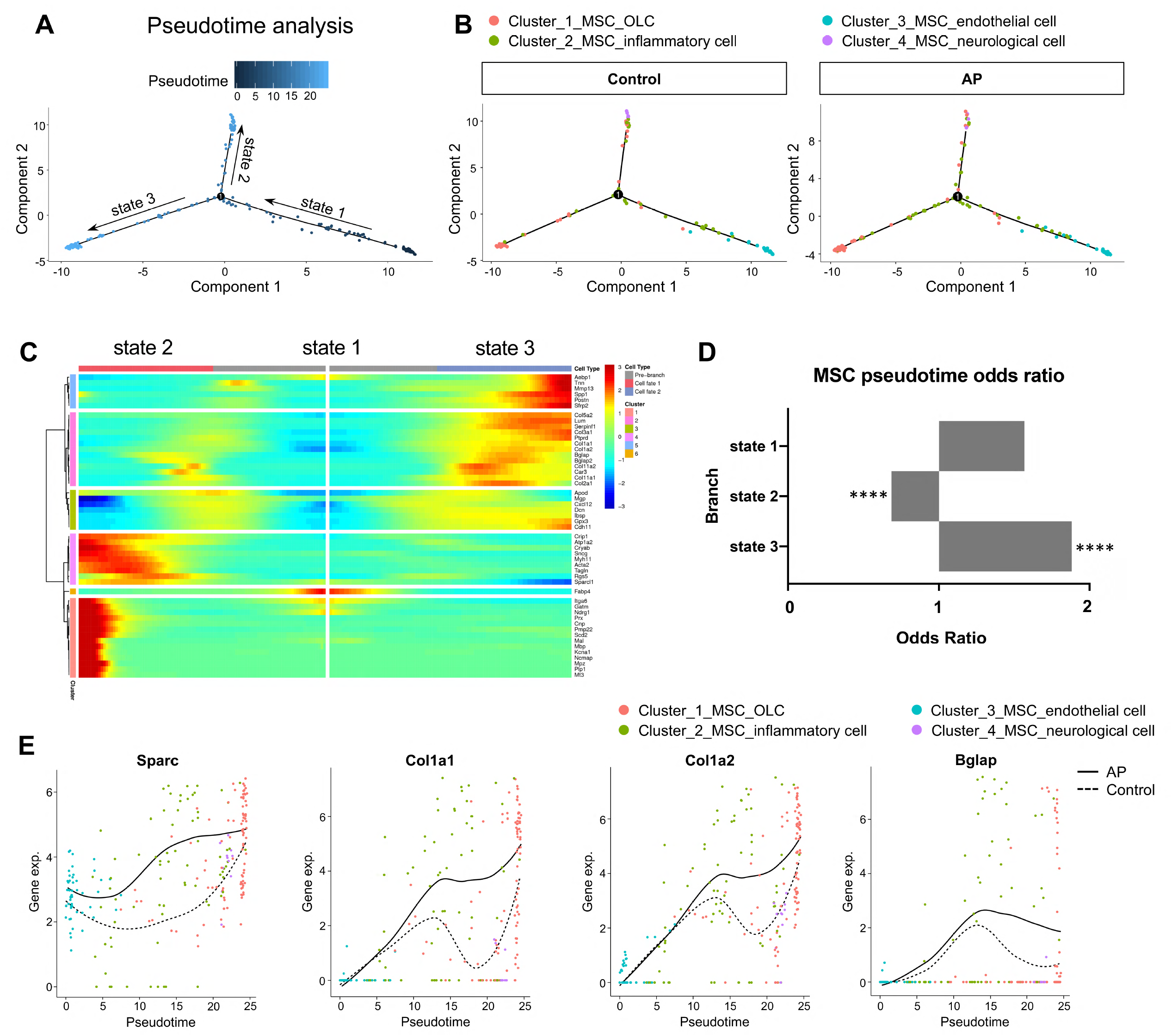
AP stimulates MSC differentiation towards osteoblast lineage cells. (A) Pseudotime analysis of the MSC population. (B) Reconstructed cell differentiation trajectory of MSC populations in control and AP groups. (C) Heatmap of differential genes of three states. (D) Bar plot of numerical changes between AP and control in respective states as identified in pseudotime analysis. Fisher’s exact test with Bonferroni correction was used. (E) Comparison between the pseudotime gene trajectories of MSC subpopulations showed upregulation of SPARC, Col1a1, Col1a2, and Bglap. Black line indicates AP and dotted lines indicate control.

### Higher OLC-associated marker levels were identified in alveolar bone from patients with AP

We next determined whether osteogenesis within the MSC population was altered in human patients with AP. The expressions of *SPARC* and *COL3α1* were significantly enhanced in alveolar bone from patients with AP, accompanied by increases in *SPP1* and *OCN* (Figure 6A). Moreover, immunostaining detected a more extensive distribution of SPARC and OCN in AP alveolar bone marrow, confirming higher osteogenic potential under AP conditions (Figure 6B, C). Upregulation of OLC-associated markers in AP lesions from patients is consistent with our previous results, verifying an augmentation of osteogenic characteristics within the MSC subcluster in alveolar bone marrow under AP conditions.

**Figure 6.**
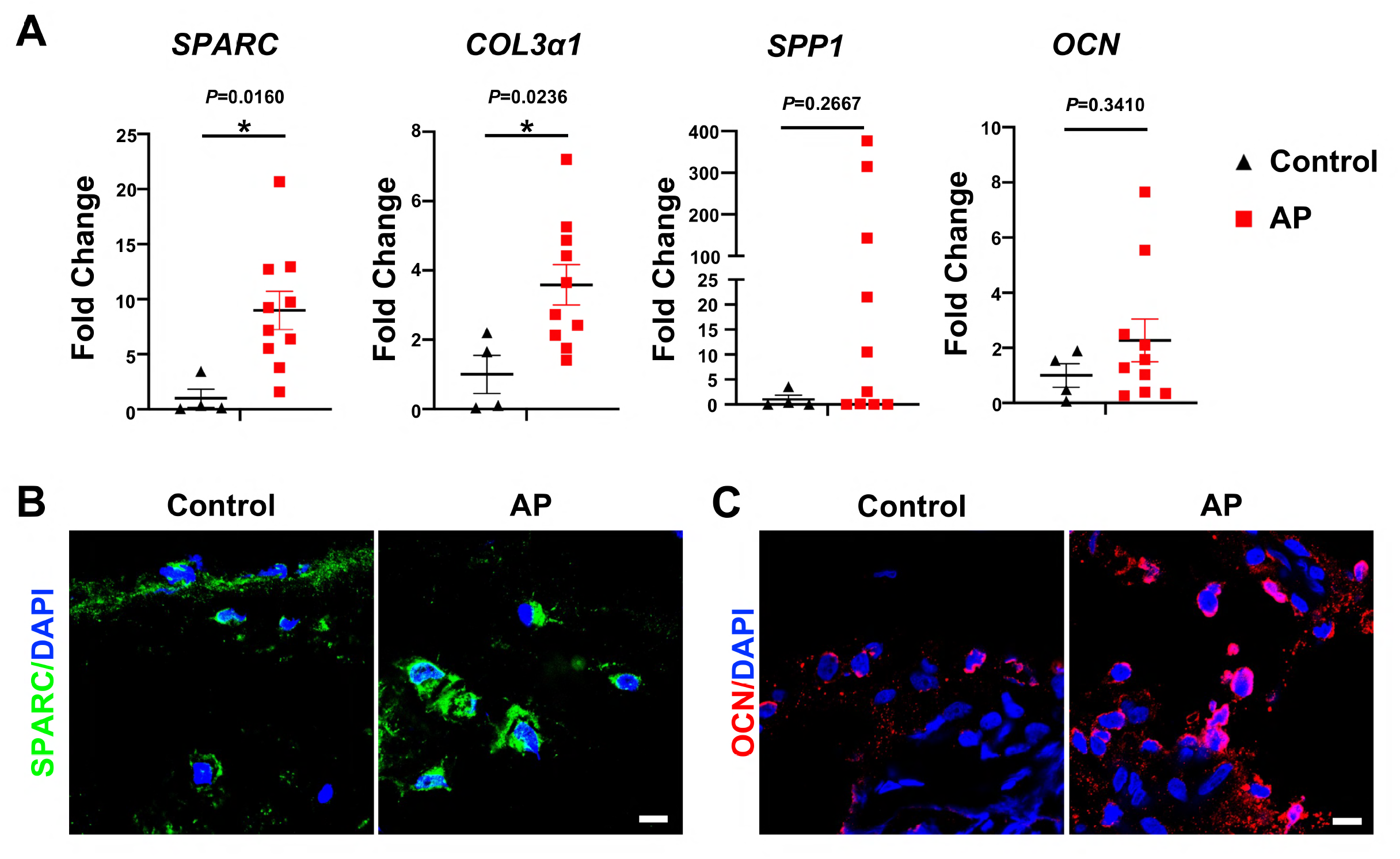
Human alveolar bone in apical periodontitis is associated with higher OLC-associated transcripts. (A) Gene expression of *SPARC, COL3α1, SPP1* and *OCN* in samples of healthy individuals and AP patients showed an upregulated expression in AP group. n=4 in healthy individuals and n=10 in AP patients. All data are shown as the mean ± SEM. (B-C) Immunofluorescence staining of SPARC and OCN in bone marrow of human alveolar bone exhibited high osteogenic protein levels in AP. n=4. Scale bar, 10 μm.

In summary, the current study elucidated the characterization of MSCs and immunoresponsive cells under healthy and chronic AP conditions, including unforeseen heterogeneity in their states of activation. We believe that this analysis presents a comprehensive transcriptomic view of the alveolar bone microenvironment during AP. These results could pave the way for designing new therapeutic approaches by targeting MSCs to restore the alveolar bone lesions caused by AP and other inflammatory-related bone diseases.

## Discussion

In this study, we employed scRNA-seq technology for an unbiased assessment of cell populations in alveolar bone marrow from mice with induced chronic apical periodontitis. We identified 15 discrete cell clusters, each with unique transcriptional signatures and putative functions during AP progression. Immune cells accounted for the majority of the cell components in the mandibular microenvironment. Cell-to-cell communication analysis revealed that among the multiple cell types, the MSC population had the most interaction with itself and other immune cells under inflammatory conditions. Furthermore, we identified four main subclusters within the MSC population. The transcriptional profiles of each subcluster from both the homeostatic and disease states were significantly different. We used the expression profile and the pseudotime analysis to demonstrate that MSCs were stimulated by inflammation and differentiated towards osteogenic lineage cells in order to protect the alveolar bone. We confirmed the enriched genes and proteins associated with osteogenesis in AP-associated alveolar bone from both the murine models and human patients. Our results define the bone marrow microenvironment in a homeostatic state and under chronic AP, highlighting the protective action of MSCs in regeneration of alveolar bone lesions.

The oral cavity is one of the most complex micro-environments where host and microbes interacts closely (Hasturk, Kantarci, & Van Dyke, 2012). Oral inflammatory diseases such as apical periodontitis is a complex disease characterized by the simultaneous presence of inflammation, destructive immunoreactions, and healing processes (Marton & Kiss, 2000; Márton & Kiss, 2014). A dense infiltration of immunocompetent cells is present in periradicular lesions. Extensive studies have examined these cells by immunostaining and gene expression techniques to determine their specific properties in terms of numbers, morphology, and function (Graunaite et al., 2012). These results provide a deeper understanding of immune cells and their inflammatory mediators. Yet the extent of their heterogeneity and the distinct markers of cell subsets have remained unexplored.

Our scRNA-seq analysis supplements knowledge of the landscape of bone marrow immunoreactions during oral inflammatory diseases. For example, the pro-inflammatory cytokines (IL-1β, IL-6, and TNF-α) have been recognized as key mediators associated with the persistence of the disease (Morsani et al., 2011). We found high expression of IL-1β and IL-6 in the mast cell cluster and TNF-α in the monocyte population. The release of these substances is directly related to both inflammatory events and bone resorption (Bracks, Armada, Goncalves, & Pires, 2014). Moreover, other inflammatory cells, including neutrophils, macrophages and lymphocytes, present specific expression patterns of various cytokines and chemokines. In particular, a significant upregulation of Cxcl2 expression was identified in neutrophil and macrophage populations. Cxcl2 was detected in endodontic infections (Gomes & Herrera, 2018) and is one of the most critical chemokines for neutrophil recruitment at a site of inflammation (Kobayashi, 2008; Silva et al., 2007). Increased levels of Cxcl2 have been reported to be associated with bone-resorptive activity in periapical lesions (AlShwaimi et al., 2013), highlighting the pro-inflammatory role of Cxcl2 in the progression of AP. Furthermore, we detected several new anti-inflammatory factors such as Ifitm1 and Ifitm2 in monocyte/macrophage subsets. Ifitm1 and Ifitm2 encode for a family of small transmembrane proteins localized in the plasma and endolysosomal membranes. Although the function of Ifitm has not been assessed in the context of AP, there is considerable evidence suggesting that Ifitm1 and Ifitm2 can be stimulated by interferons (IFNs) and exert an anti-inflammatory function in innate and adaptive immunity (Y. Liao et al., 2019; Yanez et al., 2020). Our scRNA-seq analysis has revealed novel transcriptional signatures of specific cell populations which were not visible with bulk analysis. Further studies will be required to investigate the contribution of these factors to the immune response and inflammation.

Cell-cell communications mediated by ligand-receptor complexes are crucial to control diverse biological processes, including development, differentiation and inflammation (Efremova et al., 2020). We employed CellPhoneDB to demonstrate previously unrecognized intercellular communication among all cell subsets, including close crosstalk among MSCs, macrophages and dendritic cells. More importantly, CrossTalkeR enabled us to identify cell communication differences between two biological states: disease vs. homeostasis (Nagai, Leimkuhler, et al., 2021). Our results suggest that most ligand-receptor interactions were markedly reduced during AP. Alterations in cell-cell communication have also been described for multiple tissues and organs under other disease, injury and infection conditions (Armingol, Officer, Harismendy, & Lewis, 2021). The shifts in the pattern of cell-cell interactions in the alveolar bone microenvironment may imply reorganization of the network in response to inflammation. It is worth noting that MSCs possessed the highest number of interactions with themselves and other cell types under both homeostasis and AP conditions, indicating MSCs act as a hub in cell-cell interactions. Thus, alterations of MSCs may substantially affect the biological behaviors of other cell types in both healthy and diseased states, further underscoring the importance of MSCs in alveolar bone marrow during inflammation.

In most cases of chronic AP, periapical bone destruction is self-limiting in which a new equilibrium has been established between root canal pathogens and anti-infective defense mechanisms (L. Wang, Zhang, Xiong, & Peng, 2011). The mechanisms that underly the protective actions of immune cells in apical inflammatory tissues have been extensively studied (Marton & Kiss, 2000; Márton & Kiss, 2014; Nair, 2004; Stashenko, 1990). Yet, the kinetics for the healing of these lesions by regenerating alveolar bone is less well understood.

MSCs are recognized as a promising resource for periapical bone healing in the treatment of oral inflammatory diseases. Several lines of evidence support this tenet. The evoked-bleeding step in revascularization procedures for regenerative endodontic treatment leads to a substantial influx of MSCs, confirming the existence and regenerative properties of MSCs in the periapical region. Indeed, several dental MSCs are present in the periapical tissues, including MSCs derived from alveolar bone, periodontal ligament and apical papilla (Zhao & Chai, 2015). More importantly, MSCs were found to be present in inflamed periapical tissues in adult patients with AP. Evidence showed that human apical papilla and its resident cells from teeth with pulp necrosis and AP could survive and retain their stemness in an inflammatory environment (Chrepa, Pitcher, Henry, & Diogenes, 2017). MSCs found in the periapical region were positive for typical stem cell markers, such as STRO-1, CD90, CD73, CD105, CD146 and Sox2 (Chrepa, Henry, Daniel, & Diogenes, 2015; Estrela et al., 2017; J. Liao et al., 2011). These progenitors were able to form colony-forming unit-fibroblasts (CFU-Fs), illustrating a typical MSC feature. However, our understanding of the characteristics of MSCs in inflamed periapical tissues remains at an early stage. In this study, we explored the heterogeneity and expression signature of MSCs in the presence of inflammation. Several key observations were only possible by study at the single cell level. First, we identified four main subclusters in the MSC population: MSC_OLCs, MSC_endothelial cells, MSC_neurological cells and MSC_inflammatory cells. The first three clusters were consistent with previously described MSC populations in alveolar bone marrow (Lin et al., 2021). These MSC subclusters have distinct differentiation potentials and may be associated with the pathogenesis of AP. Importantly, we noted that the proportion of MSC_inflammatory cells in AP was higher and accompanied by significantly increased expression of S100a8/a9. These factors are involved with Ca^2+^ binding which is significantly increased during inflammatory processes (S. Wang et al., 2018). Recent studies have found that although MSCs do not express S100a8/a9 under healthy conditions, they do upregulate its expression in disease states (Leimkuhler et al., 2021). The higher S100a8/a9 expression in the MSC_inflammatory cell subcluster suggests that S100a8/a9 might be a candidate for detecting an inflammatory-associated MSC state in AP.

Previous studies have shown that human apical granulation tissues contain osteogenic cells (Maeda, Wada, Nakamuta, & Akamine, 2004). However, the differentiative traits of these cells and their role in inflamed periapical tissue remain unknown. An important finding in the present study is the increased osteogenic potential in the MSC population during inflammatory-associated bone lesions. Several lines of evidence support this conclusion. The overall size of the MSC population was significantly increased and was accompanied by upregulated expression of key osteogenic-related genes. Moreover, pseudotime analysis detected a specific state progression towards osteogenesis in the MSC cluster. The numerical changes in this state were statistically higher in AP samples when compared to control. MSCs in this state harbored the highest osteoblastic differentiation potential, along with upregulated expressions of Sparc, Col1α1, Col1α2, and Bglap. More importantly, we confirmed this observation in animal models and human patients with AP. By using a lineage tracing strategy, we identified increased percentages of Prx1^+^ MSC and Sp7^+^ progenitor cells with higher expression of Sparc and Col1α1 in the alveolar bone region surrounding periapical lesions. We show, for the first time, upregulated SPARC, COL3α1 and OCN in alveolar bone from human patients with AP. It is notable that Sparc was the most upregulated marker in the MSC population from AP samples. Sparc is one of the most abundant non-collagenous proteins expressed in bone. It has a critical role in osteoblast differentiation, procollagen processing and assembly as well as mineralization (Rosset & Bradshaw, 2016). SPARC mutations have been identified in patients with osteogenesis imperfecta and idiopathic osteoporosis (Delany, McMahon, Powell, Greenberg, & Kurland, 2008; Mendoza-Londono et al., 2015). Sparc null mice exhibit decreased osteoblast number and activity along with reduced bone formation rate (Delany, Kalajzic, Bradshaw, Sage, & Canalis, 2003). Recent research using scRNA-seq revealed high levels of SPARC expression in human periodontal MSCs, highlighting the potential role of SPARC in oral MSC populations (Pagella, de Vargas Roditi, Stadlinger, Moor, & Mitsiadis, 2021). Notably, Sparc is also a molecule closely related to inflammation. Overexpressed Sparc is observed in inflammatory pulp cells, experimental glomerulonephritis and ovarian cancer associated inflammation (Dimitrova-Nakov, Baudry, Harichane, Kellermann, & Goldberg, 2014; Pichler et al., 1996; Said, Elmarakby, Imig, Fulton, & Motamed, 2008). Both pro-inflammatory and anti-inflammatory properties of Sparc have been reported (Said et al., 2008; Sangaletti et al., 2011; Tanaka et al., 2018; Toba et al., 2015). Sparc may act in two ways by activating different pathways depending on context. It therefore creates a microenvironment suitable for tissue remodeling and repair at different stages (Ghanemi, Yoshioka, & St-Amand, 2020). Given the upregulation of Sparc observed in AP mouse models and human patients, we believe that the MSC population was stimulated by inflammation and underwent osteogenesis with a significant function in bone matrix assembly and mineralization, thus serving as an important protective component in healing the bone lesions in inflammation.

In summary, we profiled the transcriptome of alveolar bone marrow single cells from control and AP samples, providing novel insights into the inflammatory biomarkers associated with the pathogenesis of oral inflammatory diseases. The interplay among various cell populations highlights the role of MSC subsets and the therapeutic potential of MSCs in healing bone lesions. These results demonstrate marked heterogeneity in the alveolar bone marrow microenvironment and provide a basis for considering MSC-based treatment for inflammatory-related bone diseases.

## Materials and methods

### Animal experiments

*Sp7Cre, Prx1Cre*, and *B6*.*Cg-Gt(ROSA)26Sortm14(CAG-tdTomato)Hze/J* mice (*Tm*^*fl/fl*^) were purchased from Jackson Laboratory. Hybrid mice were generated to mark *Sp7-* and *Prx1-*expressing cells by crossing *Sp7Cre and Prx1Cre* with *Tm*^*fl/fl*^ respectively. Wild-type mice were purchased from Chengdu Dossy Biological Technology Co., Ltd. All animal experiments were carried out in accordance with the guidelines of the Institutional Animal Care and Use Committee at the State Key Laboratory of Oral Diseases, Sichuan University (WCHSIRB-D-2021-339).

### Apical periodontitis mouse model

The apical periodontitis model was generated using eight-week-old mice as described previously (Taira et al., 2019). In brief, the pulp chambers of the mandibular first molars were opened by a high-speed handpiece with #1/4 carbide round bur under direct visualization by a stereoscopic microscope (Leica). A #10 endodontic K file was then used to probe the root canals. The teeth were then exposed to the oral environment for three weeks.

### Single cell sample preparation and sequencing

Twenty C57/B6 male mice with apical periodontitis or control mice were used to acquire single-cell suspensions. Mandibles were dissected under a stereo microscope (Leica). Specifically, soft tissues, molars, incisors and bone from behind the condyle were removed. Then, mandibular alveolar bone tissue was cut into small pieces (<1 mm^3^) and digested with 3 mg/mL collagenase type I (Gibco) and 4 g/mL dispase II (Sigma) for 60 min at 37°C as previously described (Cui et al., 2020; Yamaza et al., 2011). Red blood cells were lysed with 1 mL Red Cell Lysis Solution (Biosharp). Cells were centrifuged and resuspended in 1% BSA solution. The final concentration of cells was 1 × 10^6^ cells per mL. Cellular suspensions were loaded into Chromium microfluidic chips and barcoded with a 10x Chromium Controller (10x Genomics). RNA from the barcoded cells was subsequently reverse-transcribed and sequencing libraries constructed with reagents from a Chromium Single Cell 3’ v3 reagent kit (10x Genomics) and sequenced with the NovaSeq system (Illumina).

### Preprocessing of scRNA-seq data

We used Cellranger (v3.1, 10x Genomics) to compare reads to the mouse genome mm10 and for detection of cells using default parameters. Next, we used the Seurat package (v3.1) for further analysis of the scRNA-seq. A gene with less than 3 cells, mitochondrial genes higher than 20%, cells with less than 200 expressed genes and cells with erythrocyte gene expression higher than 5% were filtered out. The Seurat package was used for data normalization and dimensional reduction. Principal component analysis (PCA) was based on the highly variable genes, and subclusters of cells were identified using specific gene expression in each group.

### Pseudotime analysis

The cluster identified as MSC was loaded into R environment, and then the R package Monocle 2 was used to infer the trajectory and reduce the dimensional space to discover the state transitions of MSCs (Qiu, Mao, et al., 2017). Each cell’s pseudotime was automatically calculated and the branch was assigned on the principal graph. Branch expression analysis modelling (BEAM) was run to analyze the hub genes in branches and top 50 genes were visualized with the plot_genes_branched_heatmap function.

### Cell-cell interaction analysis

CellPhoneDB v.2.0 was performed to predict enriched cellular interaction between different cell populations according to the expression of a receptor by one state and a ligand by another one (Efremova et al., 2020). Receptors and ligands were analyzed when they were expressed in at least 10% cells of the corresponding cluster in mouse data. Subsequently, CrossTalkeR was used to visualize the ligand-receptor networks, which identified relevant ligands, receptors and cell types contributing to changes in cellular crosstalk when contrasting AP and homeostasis states (Nagai, Leimkühler, Schaub, Schneider, & Costa, 2021). We then ranked the individual ligand by their number of interactions and normalized to the number of all possible cell-cell interactions to identify the most influential ligand. Interactions associated with Fn1 and Sele were visualized via Sankey plot.

### Collection of human alveolar bone

This study was permitted by the Ethical Committees of the West China Hospital of Stomatology, Sichuan University (WCHSIRB-D-2021-292). Written informed consent was obtained from all patients. Patients who were characterized with periapical rarefaction in radiography, suffered with a failed root canal treatment and needed endodontic surgery were recruited as subjects. Patients with periodontitis, vertical root fracture, suspected odontogenic tumor, systematic diseases, or a history of antibiotic, antiviral or immunosuppressive therapy for 3 months were excluded. Ten human alveolar bone samples were collected from discards during the apicoectomy surgery. Normal apical bone tissue was harvested from patients who required a bone repair procedure. There was no significant difference in age or gender between the control group and the patients with apical periodontitis. The samples were rinsed by sterile PBS. Subsequently, part of the samples was quick-frozen with liquid nitrogen and stored at −80°C for mRNA extraction. Others were fixed by 4% PFA and decalcified by 20% EDTA for immunostaining.

### Quantitative Real-time PCR (qRT-PCR)

PowerLyzer 24 Homogenizer (Qiagen) was used for lysis and homogenization of alveolar bone samples and Trizol (Invitrogen) was used for mRNA extraction according to the protocol. NanoDrop ND-1000 (Thermo fisher Scientific) was used to quantify RNA concentration. PrimeScript RT reagent kit (Takara) reverse transcribed RNA and qPT-PCR was performed by SybrGreen Supermix (Bio-Rad Laboratories), with β-actin for normalization. All primers are listed in the table 1.

### Immunostaining

Immunofluorescent staining was performed to assess the expression of proteins in the region of the apical periodontitis lesion. Mandibles from *Sp7Cre;Tm*^*fl/+*^ *and Prx1Cre;Tm*^*fl/+*^ mice were dissected and fixed in 4% paraformaldehyde for three days. Subsequently, samples were decalcified in 20% EDTA (pH 7.5) and embedded in Tissue-Tek O.C.T Compound (Sakura) and cut into 8 μm sections using CryoStar NX50 (Thermo fisher Scientific). Slides were permeabilized with 0.3% Triton X-100 (Beyotime) for 15 min, blocked with 5% BSA for 1 h, incubated with anti-Sparc (1:100, R&D, AF942) or anti-Collagen I (1:200, Abcam, ab21286) or anti-Ocn (1:200, Abcam, ab93876) overnight at 4 °C, and then incubated with Alexa fluor 488 (1:1000, Invitrogen, A11070) for 1 h at room temperature. DAPI (Vector) was used as a nuclei counterstain. An Olympus confocal microscope FV3000 (Olympus) was used to capture images.

### Statistical analysis

GraphPad Prism 9.0 (GraphPad Software Inc.) was used for statistical analysis. Unpaired two-tailed Student’s t test was used in two-group comparisons. All data were exhibited as mean ± SEM. *P* values <0.05 were considered statistically significant for all analyses.

## Acknowledgements

This work was supported by NSFC grants 81800928, 81901040, and 82171001, the Young Elite Scientist Sponsorship Program by CAST (No. 2020QNRC001 and 2018QNR001), the Sichuan Science and Technology Program (No. 2019YJ0054), Research Funding from West China School/Hospital of Stomatology Sichuan University (No. RCDWJS2021-1), State Key Laboratory of Oral Diseases Open Funding Grant SKLOD-R013.

## Data availability

All data generated or analyzed during this study are included in the manuscript, supporting file and source data files. Sequence data are deposited in the NCBI Gene Expression Omnibus under accession numbers GSE212975.

## Additional information

### Author contributions

C.Z., Q.Y., and Y.F. conceived the study and designed all experiments. Y.F., P.L, C.C., C.Z., Q.Y., and C.J.R. wrote the manuscript. Y.F., P.L. C.C., R.X., and R.B. performed the experiments and analyzed data. All authors have read and approved the article.

## Figure legend

**Figure 1-figure supplement 1.**
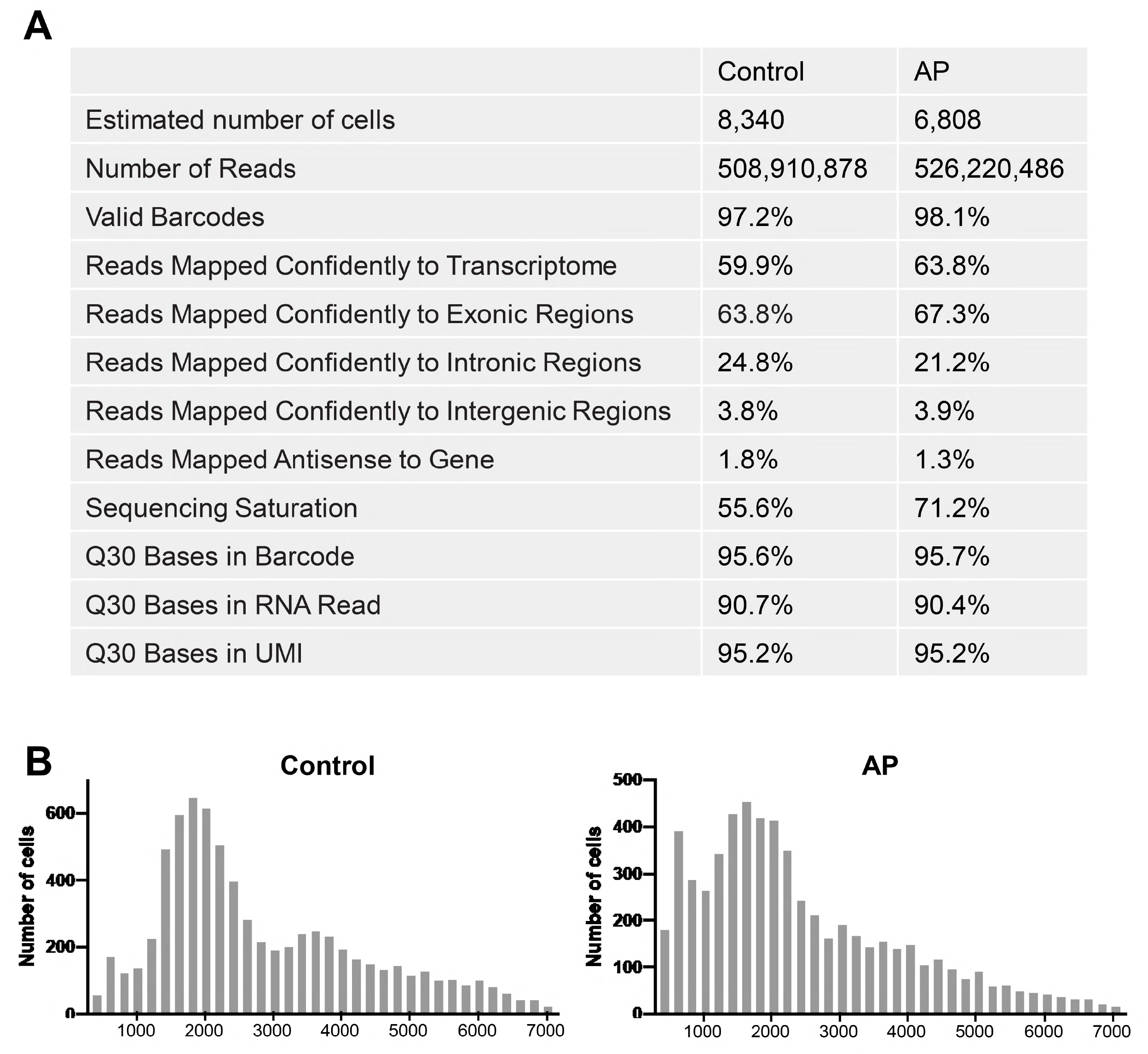
Quality control results of the sequencing data.

**Figure 4-figure supplement 1.**
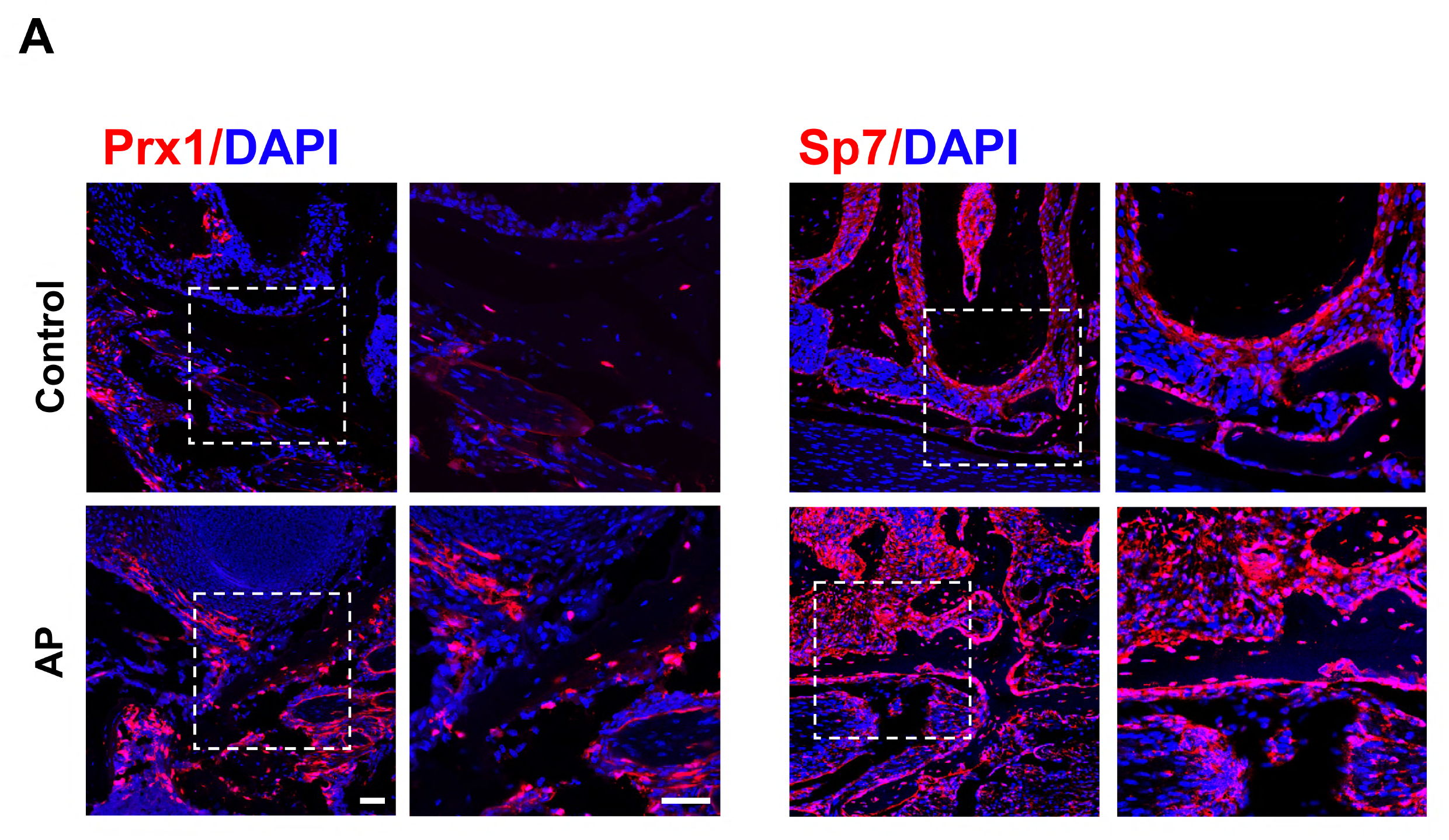
TdTomato expression in *Prx1Cre;Tm*^*fl/+*^ and *Sp7Cre;Tm*^*fl/+*^ mouse models.

**Figure 4-figure supplement 2.**
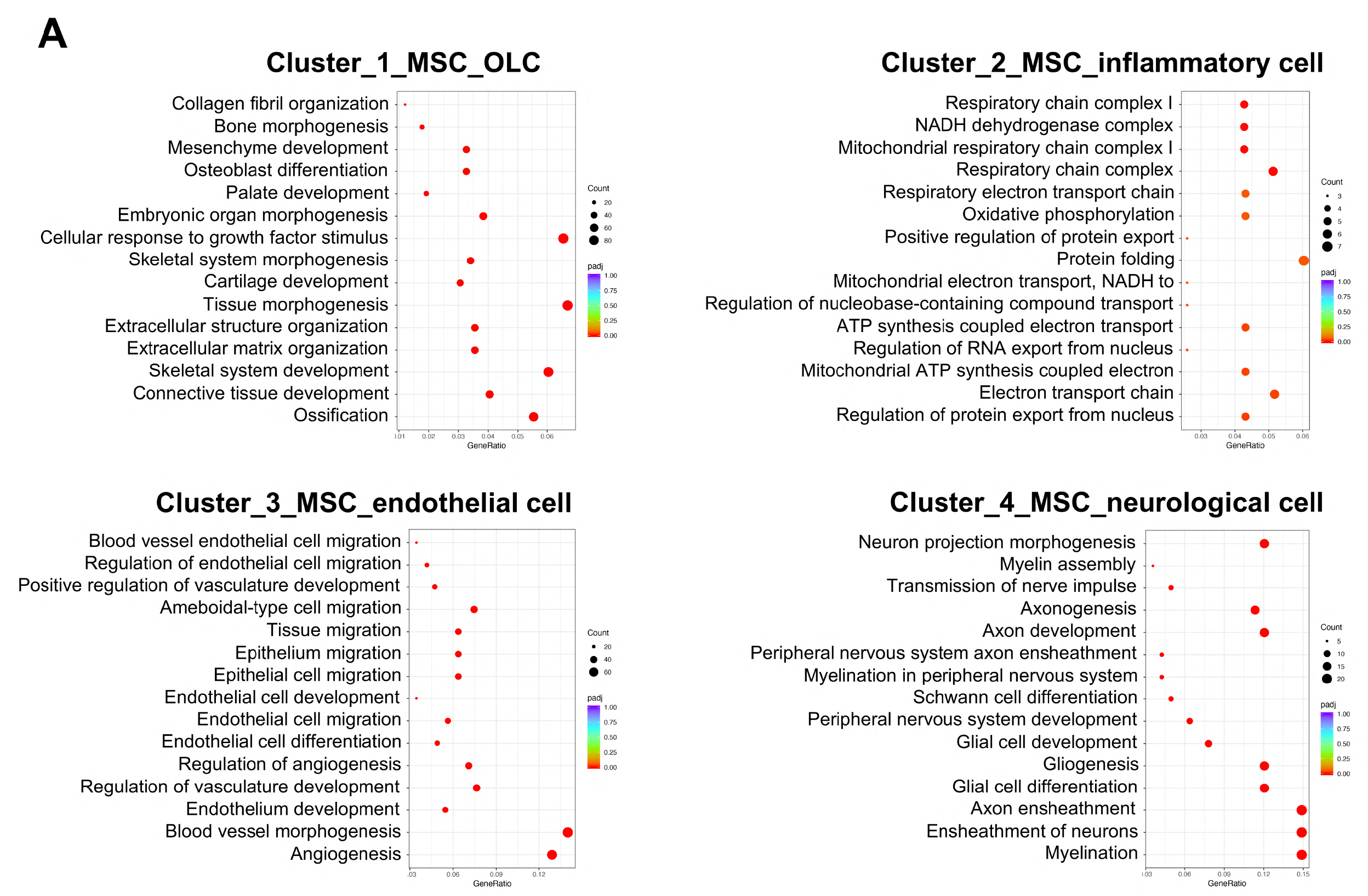
GO enrichment analysis of the biological functions of MSC subclusters.

**Supplemental Table 1.**
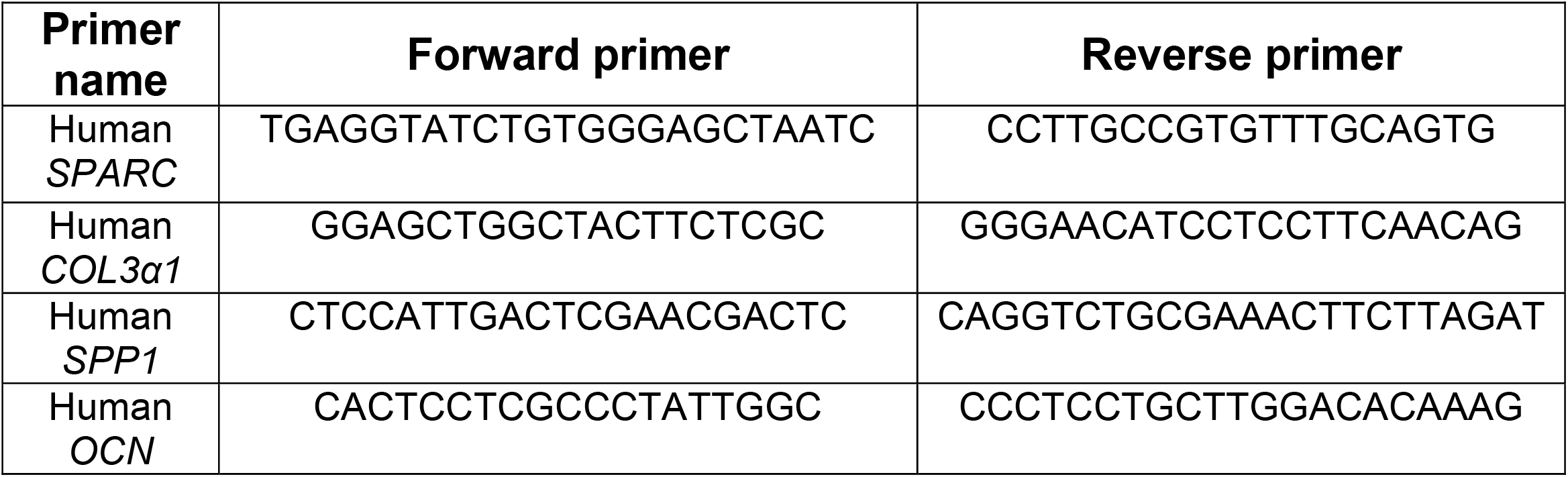
qRT-PCR primer sequences.

## References

AlShwaimi, E., Berggreen, E., Furusho, H., Rossall, J. C., Dobeck, J., Yoganathan, S., … Sasaki, H. (2013). IL-17 receptor A signaling is protective in infection-stimulated periapical bone destruction. J Immunol, 191(4), 1785–1791. doi:10.4049/jimmunol.1202194

Araujo-Pires, A. C., Biguetti, C. C., Repeke, C. E., Rodini Cde, O., Campanelli, A. P., Trombone, A. P., … Garlet, G. P. (2014). Mesenchymal stem cells as active prohealing and immunosuppressive agents in periapical environment: evidence from human and experimental periapical lesions. J Endod, 40(10), 1560–1565. doi:10.1016/j.joen.2014.02.012

Armingol, E., Officer, A., Harismendy, O., & Lewis, N. E. (2021). Deciphering cell-cell interactions and communication from gene expression. Nat Rev Genet, 22(2), 71–88. doi:10.1038/s41576-020-00292-x

Bracks, I. V., Armada, L., Goncalves, L. S., & Pires, F. R. (2014). Distribution of mast cells and macrophages and expression of interleukin-6 in periapical cysts. J Endod, 40(1), 63–68. doi:10.1016/j.joen.2013.09.037

Braz-Silva, P. H., Bergamini, M. L., Mardegan, A. P., De Rosa, C. S., Hasseus, B., & Jonasson, P. (2019). Inflammatory profile of chronic apical periodontitis: a literature review. Acta Odontol Scand, 77(3), 173–180. doi:10.1080/00016357.2018.1521005

Carminho-Rodrigues, M. T., Klee, P., Laurent, S., Guipponi, M., Abramowicz, M., Cao-van, H., … Paoloni-Giacobino, A. (2020). LARS2-Perrault syndrome: a new case report and literature review. BMC Med Genet, 21(1), 109. doi:10.1186/s12881-020-01028-8

Chrepa, V., Henry, M. A., Daniel, B. J., & Diogenes, A. (2015). Delivery of Apical Mesenchymal Stem Cells into Root Canals of Mature Teeth. J Dent Res, 94(12), 1653–1659. doi:10.1177/0022034515596527

Chrepa, V., Pitcher, B., Henry, M. A., & Diogenes, A. (2017). Survival of the Apical Papilla and Its Resident Stem Cells in a Case of Advanced Pulpal Necrosis and Apical Periodontitis. J Endod, 43(4), 561–567. doi:10.1016/j.joen.2016.09.024

Clatworthy, M. R., & Smith, K. G. (2004). FcgammaRIIb balances efficient pathogen clearance and the cytokine-mediated consequences of sepsis. J Exp Med, 199(5), 717–723. doi:10.1084/jem.20032197

Cotti, E., Schirru, E., Acquas, E., & Usai, P. (2014). An overview on biologic medications and their possible role in apical periodontitis. J Endod, 40(12), 1902–1911. doi:10.1016/j.joen.2014.08.013

Cui, C., Bi, R., Liu, W., Guan, S., Li, P., Song, D., … Fan, Y. (2020). Role of PTH1R Signaling in Prx1(+) Mesenchymal Progenitors during Eruption. J Dent Res, 99(11), 1296–1305. doi:10.1177/0022034520934732

Del Prete, A., Martinez-Munoz, L., Mazzon, C., Toffali, L., Sozio, F., Za, L., … Sozzani, S. (2017). The atypical receptor CCRL2 is required for CXCR2-dependent neutrophil recruitment and tissue damage. Blood, 130(10), 1223–1234. doi:10.1182/blood-2017-04-777680

Delany, A. M., Kalajzic, I., Bradshaw, A. D., Sage, E. H., & Canalis, E. (2003). Osteonectin-null mutation compromises osteoblast formation, maturation, and survival. Endocrinology, 144(6), 2588–2596. doi:10.1210/en.2002-221044

Delany, A. M., McMahon, D. J., Powell, J. S., Greenberg, D. A., & Kurland, E. S. (2008). Osteonectin/SPARC polymorphisms in Caucasian men with idiopathic osteoporosis. Osteoporos Int, 19(7), 969–978. doi:10.1007/s00198-007-0523-9

Dimitrova-Nakov, S., Baudry, A., Harichane, Y., Kellermann, O., & Goldberg, M. (2014). Pulp stem cells: implication in reparative dentin formation. J Endod, 40(4 Suppl), S13–18. doi:10.1016/j.joen.2014.01.011

Disease, G. B. D., Injury, I., & Prevalence, C. (2018). Global, regional, and national incidence, prevalence, and years lived with disability for 354 diseases and injuries for 195 countries and territories, 1990-2017: a systematic analysis for the Global Burden of Disease Study 2017. Lancet, 392(10159), 1789–1858. doi:10.1016/S0140-6736(18)32279-7

Efremova, M., Vento-Tormo, M., Teichmann, S. A., & Vento-Tormo, R. (2020). CellPhoneDB: inferring cell-cell communication from combined expression of multi-subunit ligand-receptor complexes. Nat Protoc, 15(4), 1484–1506. doi:10.1038/s41596-020-0292-x

Espeli, M., Smith, K. G., & Clatworthy, M. R. (2016). FcgammaRIIB and autoimmunity. Immunol Rev, 269(1), 194–211. doi:10.1111/imr.12368

Estrela, C., Carmo Souza, P. O., Barbosa, M. G., Aburad de Carvalhosa, A., Batista, A. C., Pinto Júnior, D. D. S., … de Freitas Silva, B. S. (2019). Mesenchymal Stem Cell Marker Expression in Periapical Abscess. J Endod, 45(6), 716–723. doi:10.1016/j.joen.2019.03.009

Estrela, C., Freitas Silva, B. S., Silva, J. A., Yamamoto-Silva, F. P., Pinto-Júnior, D. D., & Gomez, R. S. (2017). Stem Cell Marker Expression in Persistent Apical Periodontitis. J Endod, 43(1), 63–68. doi:10.1016/j.joen.2016.09.002

Frenette, P. S., Subbarao, S., Mazo, I. B., von Andrian, U. H., & Wagner, D. D. (1998). Endothelial selectins and vascular cell adhesion molecule-1 promote hematopoietic progenitor homing to bone marrow. Proc Natl Acad Sci U S A, 95(24), 14423–14428. doi:10.1073/pnas.95.24.14423

Gautier, E. L., Shay, T., Miller, J., Greter, M., Jakubzick, C., Ivanov, S., … Immunological Genome, C. (2012). Gene-expression profiles and transcriptional regulatory pathways that underlie the identity and diversity of mouse tissue macrophages. Nat Immunol, 13(11), 1118–1128. doi:10.1038/ni.2419

Gazivoda, D., Dzopalic, T., Bozic, B., Tatomirovic, Z., Brkic, Z., & Colic, M. (2009). Production of proinflammatory and immunoregulatory cytokines by inflammatory cells from periapical lesions in culture. J Oral Pathol Med, 38(7), 605–611. doi:10.1111/j.1600-0714.2009.00788.x

Ghanemi, A., Yoshioka, M., & St-Amand, J. (2020). Secreted protein acidic and rich in cysteine and inflammation: Another homeostatic property? Cytokine, 133, 155179. doi:10.1016/j.cyto.2020.155179

Gomes, B., & Herrera, D. R. (2018). Etiologic role of root canal infection in apical periodontitis and its relationship with clinical symptomatology. Braz Oral Res, 32(suppl 1), e69. doi:10.1590/1807-3107bor-2018.vol32.0069

Graunaite, I., Lodiene, G., & Maciulskiene, V. (2012). Pathogenesis of apical periodontitis: a literature review. J Oral Maxillofac Res, 2(4), e1. doi:10.5037/jomr.2011.2401

Hasturk, H., Kantarci, A., & Van Dyke, T. E. (2012). Oral inflammatory diseases and systemic inflammation: role of the macrophage. Front Immunol, 3, 118. doi:10.3389/fimmu.2012.00118

Hwang, B., Lee, J. H., & Bang, D. (2018). Single-cell RNA sequencing technologies and bioinformatics pipelines. Exp Mol Med, 50(8), 1–14. doi:10.1038/s12276-018-0071-8

Italiani, P., & Boraschi, D. (2014). From Monocytes to M1/M2 Macrophages: Phenotypical vs. Functional Differentiation. Front Immunol, 5, 514. doi:10.3389/fimmu.2014.00514

Jofre-Monseny, L., Loboda, A., Wagner, A. E., Huebbe, P., Boesch-Saadatmandi, C., Jozkowicz, A., … Rimbach, G. (2007). Effects of apoE genotype on macrophage inflammation and heme oxygenase-1 expression. Biochem Biophys Res Commun, 357(1), 319–324. doi:10.1016/j.bbrc.2007.03.150

Kobayashi, Y. (2008). The role of chemokines in neutrophil biology. Front Biosci, 13, 2400–2407. doi:10.2741/2853

Kolaczkowska, E., & Kubes, P. (2013). Neutrophil recruitment and function in health and inflammation. Nat Rev Immunol, 13(3), 159–175. doi:10.1038/nri3399

Leimkuhler, N. B., Gleitz, H. F. E., Ronghui, L., Snoeren, I. A. M., Fuchs, S. N. R., Nagai, J. S., … Schneider, R. K. (2021). Heterogeneous bone-marrow stromal progenitors drive myelofibrosis via a druggable alarmin axis. Cell Stem Cell, 28(4), 637–652 e638. doi:10.1016/j.stem.2020.11.004

Li, Z., Jiang, C. M., An, S., Cheng, Q., Huang, Y. F., Wang, Y. T., … Wang, J. (2014). Immunomodulatory properties of dental tissue-derived mesenchymal stem cells. Oral Dis, 20(1), 25–34. doi:10.1111/odi.12086

Liao, J., Al Shahrani, M., Al-Habib, M., Tanaka, T., & Huang, G. T. (2011). Cells isolated from inflamed periapical tissue express mesenchymal stem cell markers and are highly osteogenic. J Endod, 37(9), 1217–1224. doi:10.1016/j.joen.2011.05.022

Liao, Y., Goraya, M. U., Yuan, X., Zhang, B., Chiu, S. H., & Chen, J. L. (2019). Functional Involvement of Interferon-Inducible Transmembrane Proteins in Antiviral Immunity. Front Microbiol, 10, 1097. doi:10.3389/fmicb.2019.01097

Lin, W., Li, Q., Zhang, D., Zhang, X., Qi, X., Wang, Q., … Yuan, Q. (2021). Mapping the immune microenvironment for mandibular alveolar bone homeostasis at single-cell resolution. Bone Res, 9(1), 17. doi:10.1038/s41413-021-00141-5

Maeda, H., Wada, N., Nakamuta, H., & Akamine, A. (2004). Human periapical granulation tissue contains osteogenic cells. Cell Tissue Res, 315(2), 203–208. doi:10.1007/s00441-003-0832-z

Mantovani, A., Cassatella, M. A., Costantini, C., & Jaillon, S. (2011). Neutrophils in the activation and regulation of innate and adaptive immunity. Nat Rev Immunol, 11(8), 519–531. doi:10.1038/nri3024

Marton, I. J., & Kiss, C. (2000). Protective and destructive immune reactions in apical periodontitis. Oral Microbiol Immunol, 15(3), 139–150. doi:10.1034/j.1399-302x.2000.150301.x

Márton, I. J., & Kiss, C. (2014). Overlapping protective and destructive regulatory pathways in apical periodontitis. J Endod, 40(2), 155–163. doi:10.1016/j.joen.2013.10.036

Mendoza-Londono, R., Fahiminiya, S., Majewski, J., CareRare Canada, C., Tetreault, M., Nadaf, J., … Rauch, F. (2015). Recessive osteogenesis imperfecta caused by missense mutations in SPARC. Am J Hum Genet, 96(6), 979–985. doi:10.1016/j.ajhg.2015.04.021

Morsani, J. M., Aminoshariae, A., Han, Y. W., Montagnese, T. A., & Mickel, A. (2011). Genetic predisposition to persistent apical periodontitis. J Endod, 37(4), 455–459. doi:10.1016/j.joen.2011.01.009

Nagai, J. S., Leimkuhler, N. B., Schaub, M. T., Schneider, R. K., & Costa, I. G. (2021). CrossTalkeR: Analysis and Visualisation of Ligand Receptor Networks. Bioinformatics. doi:10.1093/bioinformatics/btab370

Nagai, J. S., Leimkühler, N. B., Schaub, M. T., Schneider, R. K., & Costa, I. G. (2021). CrossTalkeR: Analysis and Visualisation of Ligand Receptor Networks. Bioinformatics. doi:10.1093/bioinformatics/btab370

Nair, P. N. (2004). Pathogenesis of apical periodontitis and the causes of endodontic failures. Crit Rev Oral Biol Med, 15(6), 348–381. doi:10.1177/154411130401500604

Ng, Y. L., Mann, V., Rahbaran, S., Lewsey, J., & Gulabivala, K. (2008). Outcome of primary root canal treatment: systematic review of the literature -- Part 2. Influence of clinical factors. Int Endod J, 41(1), 6–31. doi:10.1111/j.1365-2591.2007.01323.x

Pagella, P., de Vargas Roditi, L., Stadlinger, B., Moor, A. E., & Mitsiadis, T. A. (2021). A single-cell atlas of human teeth. iScience, 24(5), 102405. doi:10.1016/j.isci.2021.102405

Pichler, R. H., Bassuk, J. A., Hugo, C., Reed, M. J., Eng, E., Gordon, K. L., … Johnson, R. J. (1996). SPARC is expressed by mesangial cells in experimental mesangial proliferative nephritis and inhibits platelet-derived-growth-factor-medicated mesangial cell proliferation in vitro. Am J Pathol, 148(4), 1153–1167.

Qiu, X., Hill, A., Packer, J., Lin, D., Ma, Y. A., & Trapnell, C. (2017). Single-cell mRNA quantification and differential analysis with Census. Nat Methods, 14(3), 309–315. doi:10.1038/nmeth.4150

Qiu, X., Mao, Q., Tang, Y., Wang, L., Chawla, R., Pliner, H. A., & Trapnell, C. (2017). Reversed graph embedding resolves complex single-cell trajectories. Nat Methods, 14(10), 979–982. doi:10.1038/nmeth.4402

Rosset, E. M., & Bradshaw, A. D. (2016). SPARC/osteonectin in mineralized tissue. Matrix Biol, 52-54, 78–87. doi:10.1016/j.matbio.2016.02.001

Said, N. A., Elmarakby, A. A., Imig, J. D., Fulton, D. J., & Motamed, K. (2008). SPARC ameliorates ovarian cancer-associated inflammation. Neoplasia, 10(10), 1092–1104. doi:10.1593/neo.08672

Sangaletti, S., Tripodo, C., Cappetti, B., Casalini, P., Chiodoni, C., Piconese, S., … Colombo, M. P. (2011). SPARC oppositely regulates inflammation and fibrosis in bleomycin-induced lung damage. Am J Pathol, 179(6), 3000–3010. doi:10.1016/j.ajpath.2011.08.027

Shapouri-Moghaddam, A., Mohammadian, S., Vazini, H., Taghadosi, M., Esmaeili, S. A., Mardani, F., … Sahebkar, A. (2018). Macrophage plasticity, polarization, and function in health and disease. J Cell Physiol, 233(9), 6425–6440. doi:10.1002/jcp.26429

Silva, T. A., Garlet, G. P., Fukada, S. Y., Silva, J. S., & Cunha, F. Q. (2007). Chemokines in oral inflammatory diseases: apical periodontitis and periodontal disease. J Dent Res, 86(4), 306–319. doi:10.1177/154405910708600403

Stashenko, P. (1990). Role of immune cytokines in the pathogenesis of periapical lesions. Endod Dent Traumatol, 6(3), 89–96. doi:10.1111/j.1600-9657.1990.tb00400.x

Hart, L. M., Hansen, T., Rietveld, I., Dekker, J. M., Nijpels, G., Janssen, G. M., … Maassen, J. A. (2005). Evidence that the mitochondrial leucyl tRNA synthetase (LARS2) gene represents a novel type 2 diabetes susceptibility gene. Diabetes, 54(6), 1892–1895. doi:10.2337/diabetes.54.6.1892

Taira, T. M., Lima, V., Prado, D. S., Silva, T. A., Issa, J. P. M., da Silva, L. A. B., … Fukada, S. Y. (2019). NLRP12 Attenuates Inflammatory Bone Loss in Experimental Apical Periodontitis. J Dent Res, 98(4), 476–484. doi:10.1177/0022034518820289

Tanaka, M., Takagi, T., Naito, Y., Uchiyama, K., Hotta, Y., Toyokawa, Y., … Itoh, Y. (2018). Secreted protein acidic and rich in cysteine functions in colitis via IL17A regulation in mucosal CD4(+) T cells. J Gastroenterol Hepatol, 33(3), 671–680. doi:10.1111/jgh.13842

Tang, F., Barbacioru, C., Wang, Y., Nordman, E., Lee, C., Xu, N., … Surani, M. A. (2009). mRNA-Seq whole-transcriptome analysis of a single cell. Nat Methods, 6(5), 377–382. doi:10.1038/nmeth.1315

Tiburcio-Machado, C. S., Michelon, C., Zanatta, F. B., Gomes, M. S., Marin, J. A., & Bier, C. A. (2021). The global prevalence of apical periodontitis: a systematic review and meta-analysis. Int Endod J, 54(5), 712–735. doi:10.1111/iej.13467

To, W. S., & Midwood, K. S. (2011). Plasma and cellular fibronectin: distinct and independent functions during tissue repair. Fibrogenesis Tissue Repair, 4, 21. doi:10.1186/1755-1536-4-21

Toba, H., de Castro Brás, L. E., Baicu, C. F., Zile, M. R., Lindsey, M. L., & Bradshaw, A. D. (2015). Secreted protein acidic and rich in cysteine facilitates age-related cardiac inflammation and macrophage M1 polarization. Am J Physiol Cell Physiol, 308(12), C972–982. doi:10.1152/ajpcell.00402.2014

Wang, L., Zhang, R., Xiong, H., & Peng, B. (2011). The involvement of platelet-derived growth factor-A in the course of apical periodontitis. Int Endod J, 44(1), 65–71. doi:10.1111/j.1365-2591.2010.01798.x

Wang, S., Song, R., Wang, Z., Jing, Z., Wang, S., & Ma, J. (2018). S100A8/A9 in Inflammation. Front Immunol, 9, 1298. doi:10.3389/fimmu.2018.01298

Yamaza, T., Ren, G., Akiyama, K., Chen, C., Shi, Y., & Shi, S. (2011). Mouse mandible contains distinctive mesenchymal stem cells. J Dent Res, 90(3), 317–324. doi:10.1177/0022034510387796

Yanez, D. C., Ross, S., & Crompton, T. (2020). The IFITM protein family in adaptive immunity. Immunology, 159(4), 365–372. doi:10.1111/imm.13163

Zhao, H., & Chai, Y. (2015). Stem Cells in Teeth and Craniofacial Bones. J Dent Res, 94(11), 1495–1501. doi:10.1177/0022034515603972

Zheng, G. X., Terry, J. M., Belgrader, P., Ryvkin, P., Bent, Z. W., Wilson, R., … Bielas, J. H. (2017). Massively parallel digital transcriptional profiling of single cells. Nat Commun, 8, 14049. doi:10.1038/ncomms14049

